# Differential splice isoforms of mouse CDK2 play functionally redundant roles during mitotic and meiotic division

**DOI:** 10.1101/2024.09.03.610967

**Authors:** Nathan Palmer, S. Zakiah A. Talib, Jin Rong Ow, Tommaso Tabaglio, Christine M.F. Goh, Li Na Zhao, Ernesto Guccione, Kui Liu, Philipp Kaldis

## Abstract

In most mammals, the cell cycle kinase; cyclin-dependent kinase 2 (CDK2) is expressed as two major isoforms due to the inclusion or exclusion of an alternatively spliced exon. The shorter CDK2 isoform: CDK2S, is expressed constitutively during the cell cycle and can be detected in many different tissue types. In contrast, the longer isoform: CDK2L, shows preferential expression in meiotically dividing cells of the germ cells and upon S-phase entry during mitotic cell division. Both CDK2L and CDK2S form heteromeric complexes with cyclins A2 and E1 *in vitro*. However, complexes comprised of each isoform differ considerably in their kinase activity towards known CDK substrates. It is currently unknown whether the long and short isoforms of CDK2 play functionally different roles *in vivo* during either mitotic and meiotic divisions as conventional knockout methodology leads to the loss of both isoforms. In this study, we find that both CDK2L and CDK2S are sufficient to support both mitotic and meiotic division when expressed in the absence of the other. This data contributes to the explanation of the apparent tolerance of the evolutionary loss of CDK2L expression in humans.

## Introduction

Alternative splicing enables multiple protein isoforms to be generated from a single transcriptional unit thus conferring added complexity to the human transcriptome and proteome [1]. Although the vast majority of predicted alternate transcripts are thought not to be translated into functional proteins, in many cases proteins arising from alternate splice isoforms have been shown to play crucial and often divergent roles in important cellular processes [2]. In this study, we investigate the function of an alternatively spliced isoform of cyclin-dependent kinase 2 (CDK2).

During mammalian cell division, cyclin-dependent kinase 2 (CDK2) phosphorylates key proteins to promote G1/S transition and DNA replication during S-phase. These actions of CDK2 are executed in association with the E– and A-type cyclins respectively [3, 4]. Despite such central cell cycle-oriented functions, CDK2 is not essential for cellular division in several mammalian cell lines [5], and *Cdk2^-/-^* mice are viable [6, 7]. The lack of proliferative defects in *Cdk2^-/-^* cells is due to functional redundancy between CDK2 and the closely related CDK1 which is essential for normal cellular division [8–10] as well as CDK4 [11]. In the absence of CDK2, CDK1 can fulfil the roles of CDK2 by binding to D-, E– and A-type cyclins [8, 12]. Unlike during mitotic division, CDK2 is essential for meiotic divisions. In meiotically dividing cells, CDK1 is unable to compensate for specific actions of CDK2. Consequentially *Cdk2^-/-^* meiocytes undergo meiotic arrest resulting in hypoplastic reproductive organs and infertility in both male and female animals [6, 7].

In most mammals, CDK2 is expressed as two distinct protein isoforms, here referred to as CDK2S (short) or CDK2L (long) [13]. The longer CDK2L transcript is generated via alternate splicing of the CDK2 gene. In mice, this leads to the inclusion of exon 6, upon translation the inclusion of this alternatively spliced exon results in a 48 amino acid insertion. This alternate splicing event leading to the expression of CDK2L has been detected in many mammals [14–16] [17, 18] and also in the monkey COS-1 cell line [13]. The human *Cdk2* gene contains seven exons, while mouse *Cdk2* contains 8 exons [13, 19]. The DNA segment encoding the ‘extra’ exon in the mouse CDK2 gene, exon 6, also exists in the human CDK2 DNA sequence within intron 5. Although the mRNA transcript encoding the long isoform of murine *Cdk2* is listed as a putative transcript in humans (XP_011536034.1), this isoform is not expressed in several human cell lines. Additionally, it is reported that CDK2L expression is not seen in human tissues such as thymus which are known to express both long and short *Cdk2* isoforms in mice [13, 17, 20]. Although not addressed in this study, an extra layer of complexity is the fact that in humans, but not in other mammals, exon 5 is alternatively spliced. This is due to the mammalian divergence of constitutive exon 5 inclusion in the mouse *Cdk2* transcript to PCBP-responsive exon 5 alternative splicing in humans [21].

The function of the 48 amino acid insert encoded by exon 6 remains uncertain as its transcribed sequence does not share any homology with previously described protein domains. While there have been few studies investigating the differences between the functions of CDK2L and CDK2S, evidence suggests that CDK2L could play a role during pachytene and may preferentially interact with SpeedyA (see Fig.1 in [22]) in addition to differences in activating and inhibitory phosphorylation [23].

**Figure 1:**
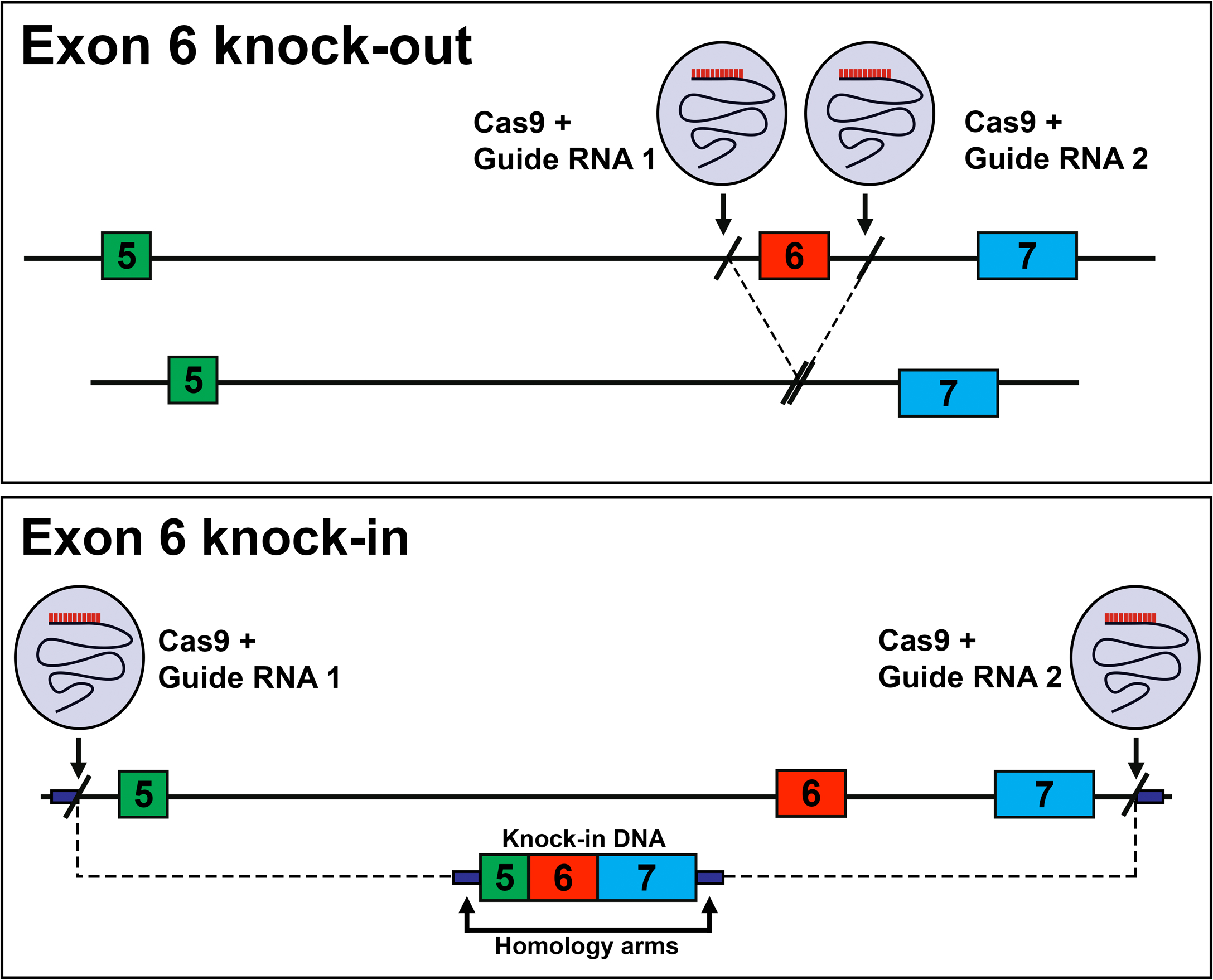
Design of the CDK2S and CDK2L mice. (upper panel) Deletion of exon 6 leads to mice that can only express CDK2S (Exon 6 knock-out; CDK2^short/short^). (lower panel) Fusion of exons 5-7 leads to the continuous expression of CDK2L and the absence of CDK2S (Exon 6 knock-in, CDK2^long/long^).

## Results

### Generation of CDK2L and CDK2S mouse models

In an attempt to identify specific biological processes mediated specifically by CDK2L or CDK2S, we generated two different mouse strains. In the first, the deletion of exon 6 completely prevents the expression of the CDK2L promoting the constitutive expression of only the shorter isoform (Figure 1; knock-out). In the second, the fusion of exons 5, 6, and 7 removes the surrounding intronic sequences required for the skipping of exon 6 forcing the sole expression of CDK2L (Figure 1; knock-in).

To confirm the correct expression of *Cdk2L* or *Cdk2S* in each of our mutant models, we performed western blotting of tissues extracted from homozygous CDK2L and CDK2S mice and compared the protein expression of CDK2 to that seen in wild type (Figure 2). CDK2L is known to be highly expressed in mouse testes and thymus [13]. In less proliferative tissues such as the brain, CDK2L cannot be detected at high levels. As previously described, CDK2 could be observed as two isoforms in wild type testis, spleen, and thymus appearing at around 39kDa (CDK2L) and 33kDa (CDK2S), but could not be detected in brain lysate (Figure 2A). In CDK2L and CDK2S mice, only the long or short isoforms could be detected, respectively (Figure 2A, lanes 5-7 and 9-11). Furthermore, these singular isoforms were expressed at levels similar to those seen for the combined expression of *Cdk2* isoforms expressed in wild type tissues, as can be seen in testis extracts (Figure 2B). Together these results suggest that CDK2L and CDK2S mouse models reported in this study correctly express only the long and short isoforms of CDK2, respectively, and that this expression occurs at similar levels to that seen for wild type CDK2 in various tissues. Phenotypically, homozygous CDK2L and CDK2S mice were found to be essentially indistinguishable from wild type mice aside from a small but significant decrease in body weight during early development.

**Figure 2:**
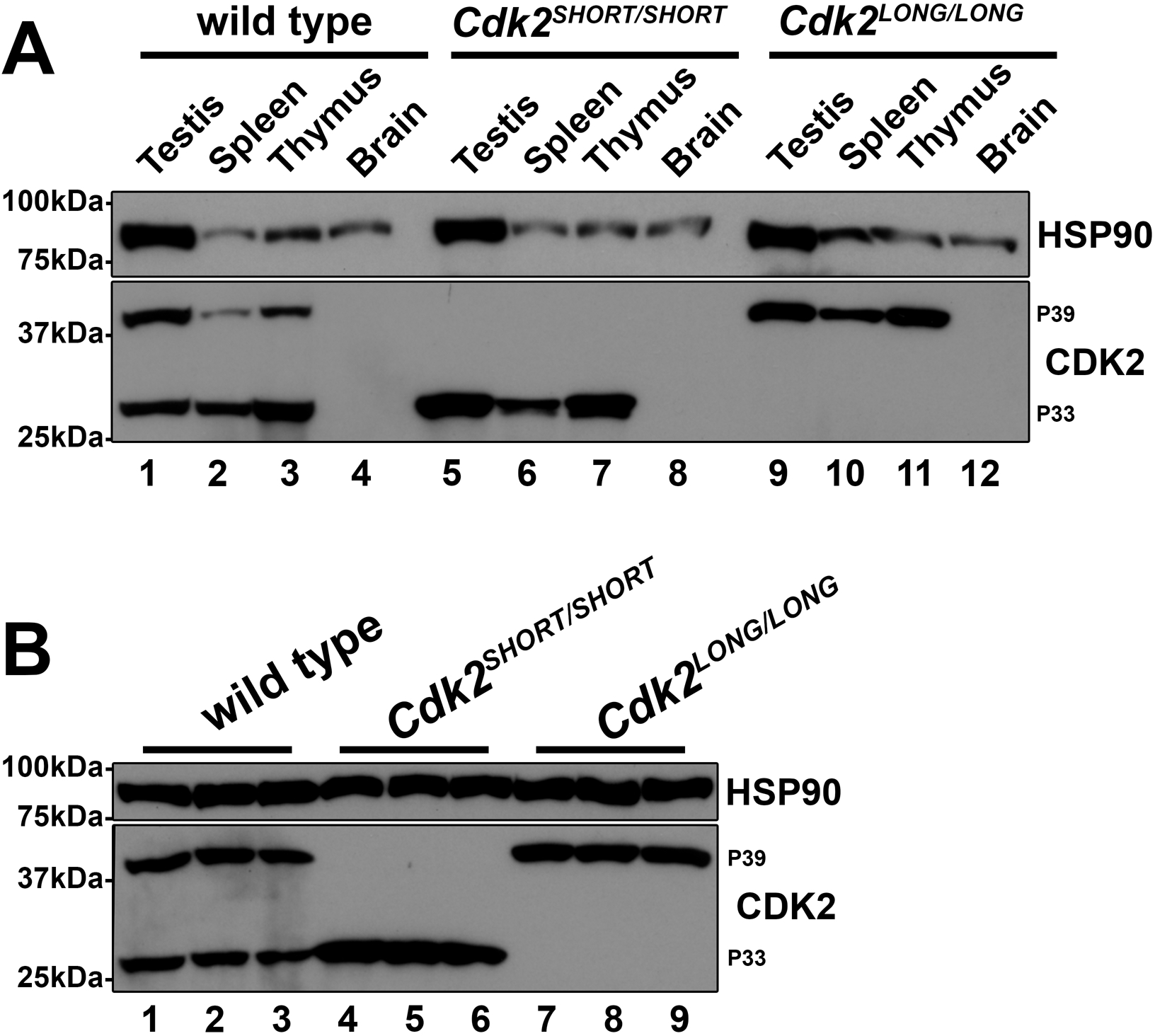
Protein expression of CDK2S and CDK2L in tissue extracts. (A) Protein extracts from testis, spleen, thymus, and brain were analyzed by western blot using pan-CDK2 antibodies. Please note that p39 corresponds to CDK2L and p33 to CDK2S, respectively. (B) Testis protein extracts from wild type, Cdk2^short/short^ (CDK2S), and Cdk2^long/long^ (CDK2L) were analyzed for CDK2 expression. Note that HSP90 serves as a loading control.

To confirm the normal development of reproductive organs, we prepared H&E stained sections from *Cdk2S* and *Cdk2L* testis and ovary. In testes of *Cdk2S* and *Cdk2L*, normal appearance of metaphase I stage primary spermatocytes, round spermatids, and elongating spermatids indicated the correct progression of spermatocytes through both meiotic divisions (Figure 3). This result was in agreement with an unchanged testis/bodyweight ratio observed in these mice (see Figure 5A). For female mice, ovarian follicles were observed to develop normally in adult CDK2L mice comparable to wild-type mice (Figure 4). Together these results demonstrate that histologically the reproductive organs of CDK2S and CDK2L mice are indistinguishable from wild type suggesting normal fertility in these models.

**Figure 3:**
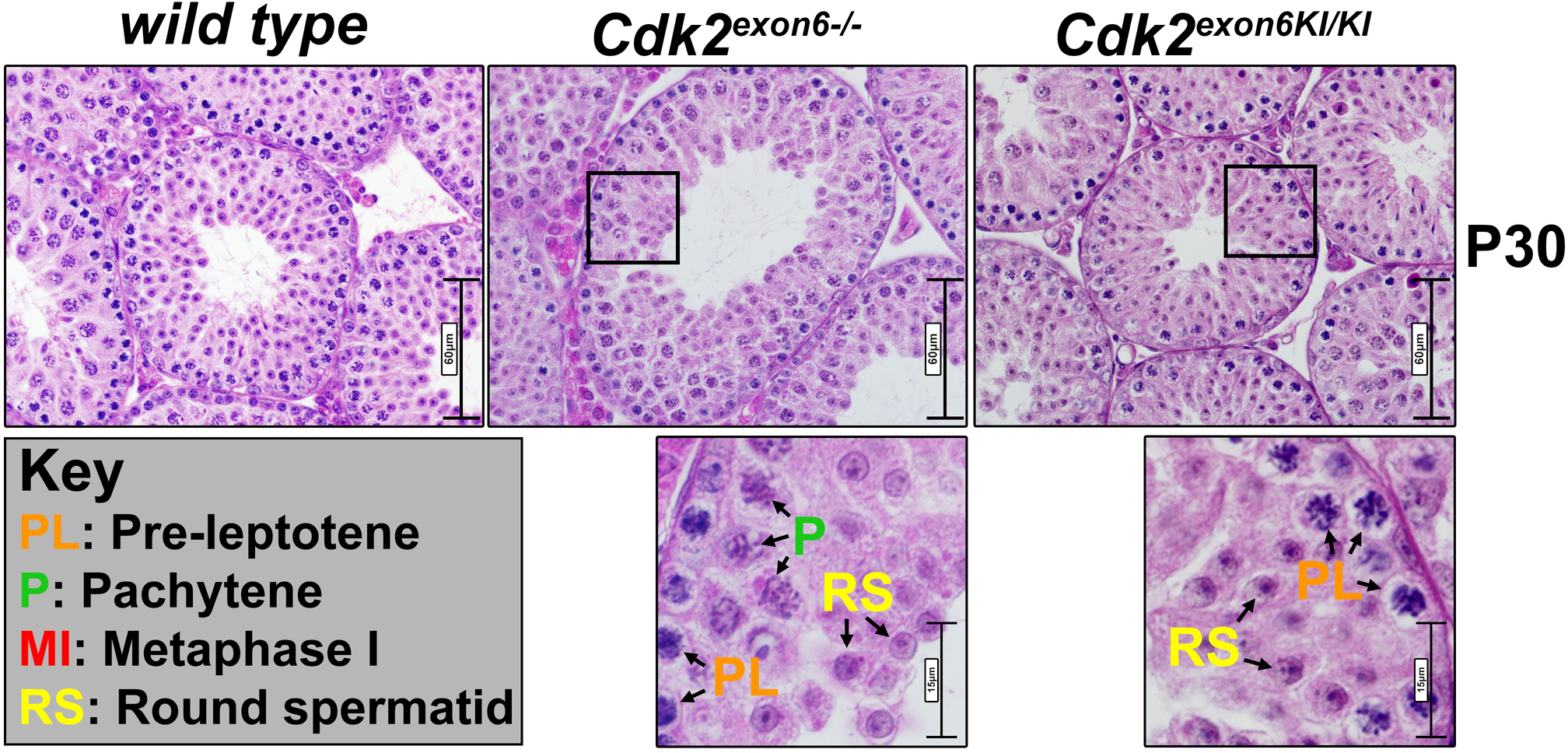
Morphology of testis. H&E stained P30 testis sections from wild type, Cdk2^short/short^ (Cdk2^exon6-/-^; CDK2S), and Cdk2^long/long^ (Cdk2^exon6KI/KI^; CDK2L) with the indicated insets. Arrows indicate pre-leptotene (PL; orange), pachytene (P; green), metaphase I (MI; red) spermatocytes, and round spermatids (RS; yellow). Scale bar is 60µm in the upper row and 15µm in the lower row.

**Figure 4:**
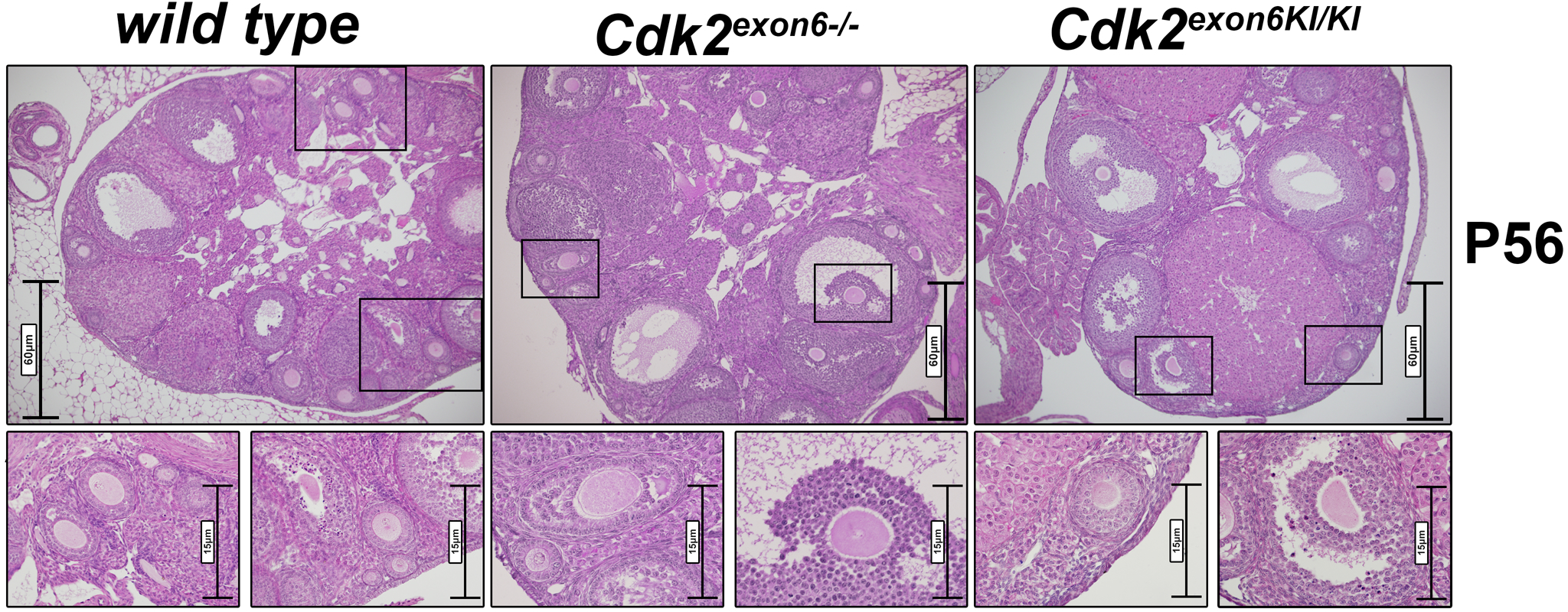
Ovaries display no obvious defects. H&E stained P56 ovary sections from wild type, Cdk2^short/short^ (Cdk2^exon6-/-^; CDK2S), and Cdk2^long/long^ (Cdk2^exon6KI/KI^; CDK2L) with the indicated insets. In the lower row, magnified follicles are depicted. Scale bar is 60µm in the upper row and 15µm in the lower row.

**Figure 5:**
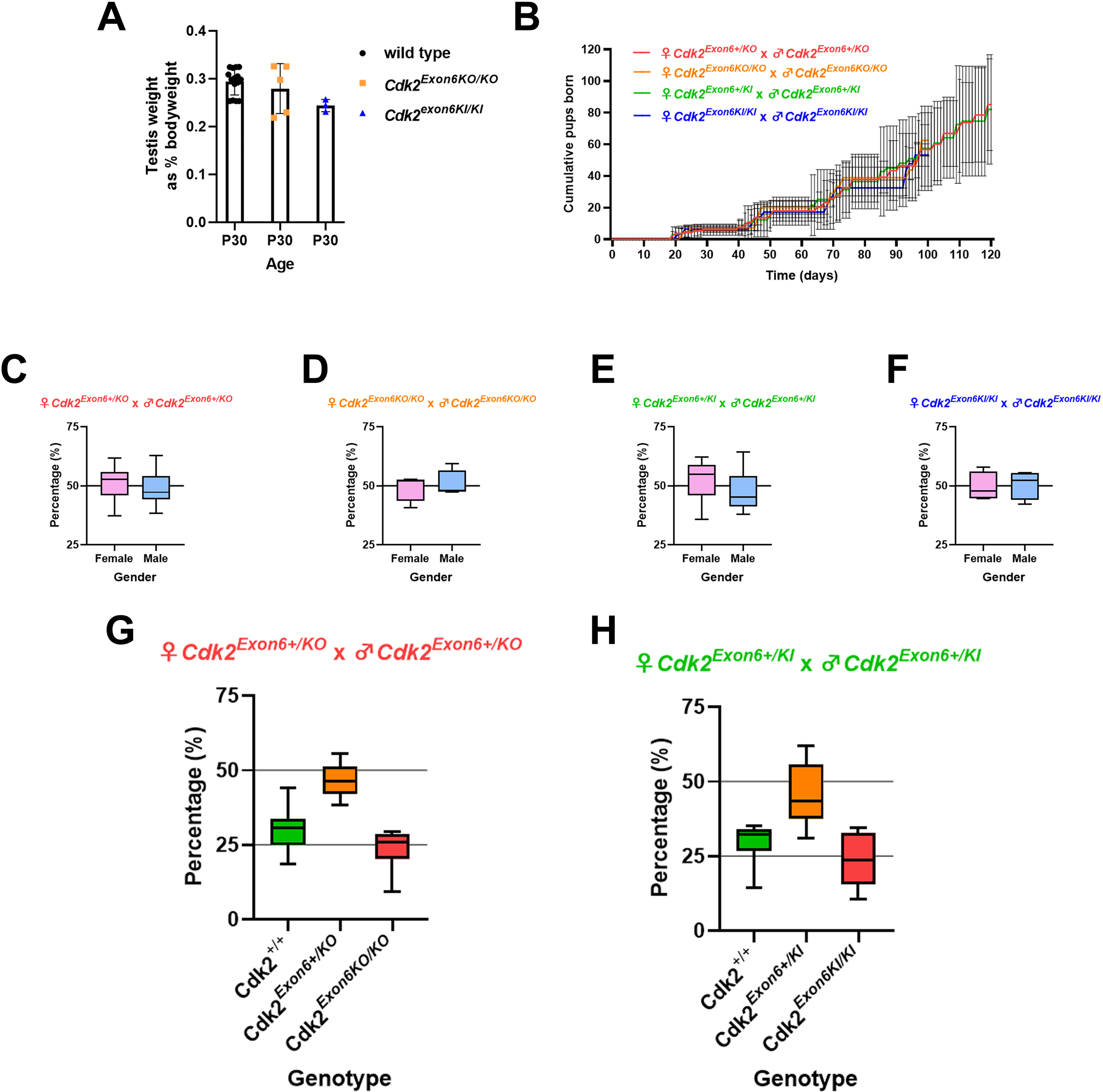
CDK2S and CDK2L mice are fertile. (A) P30 testis weight as % of bodyweight of wild type (black circles), Cdk2^short/short^ (Cdk2^exon6-/-^; CDK2S; orange squares), and Cdk2^long/long^ (Cdk2^exon6KI/KI^; CDK2L, blue triangles). (B) Heterozygote crosses of the indicated genotypes and the number of offspring (pups) during the first 120 days. (C-F) Gender balance of the pups from the breedings in (B). (G) Percentage of wild type (CDK2^+/+^), heterozygote (Cdk2^Exon6+/KO^), and homozygote (Cdk2^Exon6KO/KO^) pups from heterozygote intercrosses. (H) Percentage of wild type (CDK2^+/+^), heterozygote (Cdk2^Exon6+/KO^), and homozygote (Cdk2^Exon6KO/KO^) pups from heterozygote intercrosses. Note that the expected 25%, 50%, 25% Mendelian ratios were observed.

Remarkably, unlike *Cdk2^-/-^* mice whereby for both sexes, heterozygotes are fertile but homozygotes show sterility at 100% penetrance in crosses between adults (8-12 weeks old; [24]) heterozygous or homozygous mutant mice both led to viable pregnancies in both CDK2L and CDK2S genotypes. For each variation of the crosses performed the average litter size was not significantly different from crosses of wild type and normal Mendelian ratios were observed (Figure 5B). To determine that fertility is sustained into adulthood, we followed breedings for at least 5 litters and found that there was no obvious decrease in fecundity over this time and mutant mice continued to have comparable numbers of pups to wild type at intervals which were not significantly longer than wild type. The gender of the born pups was almost equally divided between female and male and this was not dependent on the genotype of the parents (Figure 5C-F). Finally, we analyzed heterozygote crosses and found the expected 25% (+/+), 50% (+/-), and 25% (-/-) as expected (Figure 5G-H). This data clearly shows that both CDK2S and CDK2L mice are fully fertile.

### CDK2L localizes specifically to telomeres during meiotic prophase I but can be functionally replaced by CDK2S to ensure normal homolog pairing

The normal reproductive health of CDK2L and CDK2S mutant mouse models suggests that these isoforms are functionally redundant in regards to maintaining fertility. We hypothesized that both of these isoforms must be catalytically active and able to act upon the same substrates to maintain normal meiotic division.

In mammalian meiocytes, CDK2 can be detected cytologically at telomeres throughout meiotic prophase and transiently, at late recombination nodules, specifically during mid-pachytene [25, 26]. Telomeric CDK2 is known to mediate telomeric tethering as part of the LINC (linker of nucleoplasm and cytoskeleton) complex [27, 28] (for review see [29]). In contrast, late recombination nodule-associated CDK2 orchestrates the maturation sites of meiotic recombination to form meiotic crossovers [26]. Although not currently associated with specific binding on chromosomes, further evidence also suggests that CDK2 might also act to control transcription during meiotic prophase I [30]. Upon deletion of *Cdk2* or in knock-in models where *Cdk2* is catalytically inactive, the activity of *Cdk2* at telomeres is lost. In these instances, meiotic arrest occurs as a result of the failure of homolog pairing during the zygotene-pachytene transition of meiotic prophase I [27, 28, 31]. In contrast, in knock-in models of *Cdk2* where the action of *Cdk2* at telomeres is not perturbed and homolog pairing occurs as normal [26, 32]. In this model meiotic arrest occurs due to the failure of homologous recombination between paired homologs [26].

To determine whether any meiotic defects occur following the loss of either the long or short isoforms of *Cdk2*, we prepared chromosome spread preparations from CDK2L and CDK2S spermatocytes. First, we performed dual staining for the lateral and transverse elements of the synaptonemal complex scaffolding complex SYCP3 and SYCP1. We observed normal co-localization of these proteins during meiotic prophase and the formation of 19 well-separated bivalents in addition to the XY pair indicating normal synapsis (Figure 6). As synapsis was found to be normal in CDK2L and CDK2S mice, we infer that these two isoforms of CDK2 are functionally redundant and can each act independently of the other to maintain the normal meiotic functions of CDK2.

**Figure 6:**
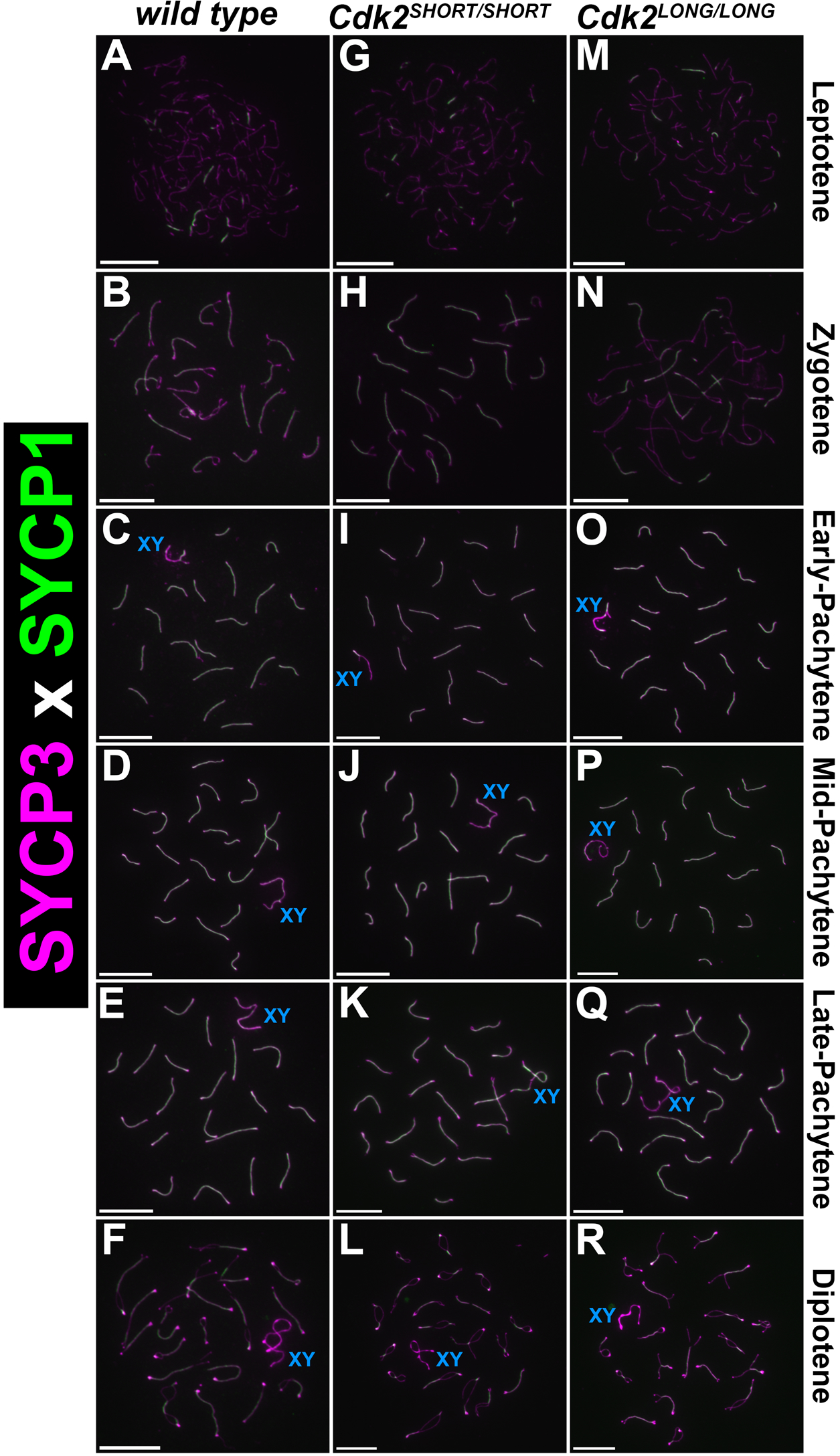
The synaptonemal complex forms normally in CDK2S and CDK2L testis. Chromosome spreads were prepared from adult testis and stained with antibodies against SYCP1 (green) and SYCP3 (cyan), respectively. Various stages of meiotic prophase I are depicted for wild type (A-F), Cdk2^short/short^ (G-L; CDK2S), and Cdk2^long/long^ (M-R; CDK2L). Spermatocytes from all 3 genotypes progress normally into mid-pachytene (D, J, P), late-pachytene (E, K, Q), and diplotene (F, L, R). The sex chromosome are labeled by a blue XY. Scale bars in all panels are 5µm. At least 3 biological replicates were analyzed for each condition.

### Repair of DNA intermediates during meiotic prophase I

During meiotic prophase I, strand invasion leads to DNA intermediates which causes apoptosis and developmental arrest in spermatocytes. A useful marker to monitor repair of DNA damage is the phosphorylated histone H2A variant (γH2AX; Ser139). The γH2AX signal in wild type spermatocytes is “cloud-like” in leptotene and zygotene (Figure 7A-B), localized along unpaired axes in early-pachytene (Figure 7C), which progressively clears up between mid-pachytene and diplotene (Figure 7D-F). The sex chromosomes remain stained by a cloud of γH2AX during all stages (Figure 7A-F). In both CDK2^SHORT/SHORT^ and CDK2^LONG/LONG^ mice that express only CDK2L and CDK2S, respectively, the pattern of γH2AX mirrored the ones in wild type mice. Therefore, we have to conclude that repair of DNA intermediates due to strand invasion is effective in the mutant mice and that early stages of meiotic prophase are not majorly affected by lacking either CDK2S or CDK2L.

**Figure 7:**
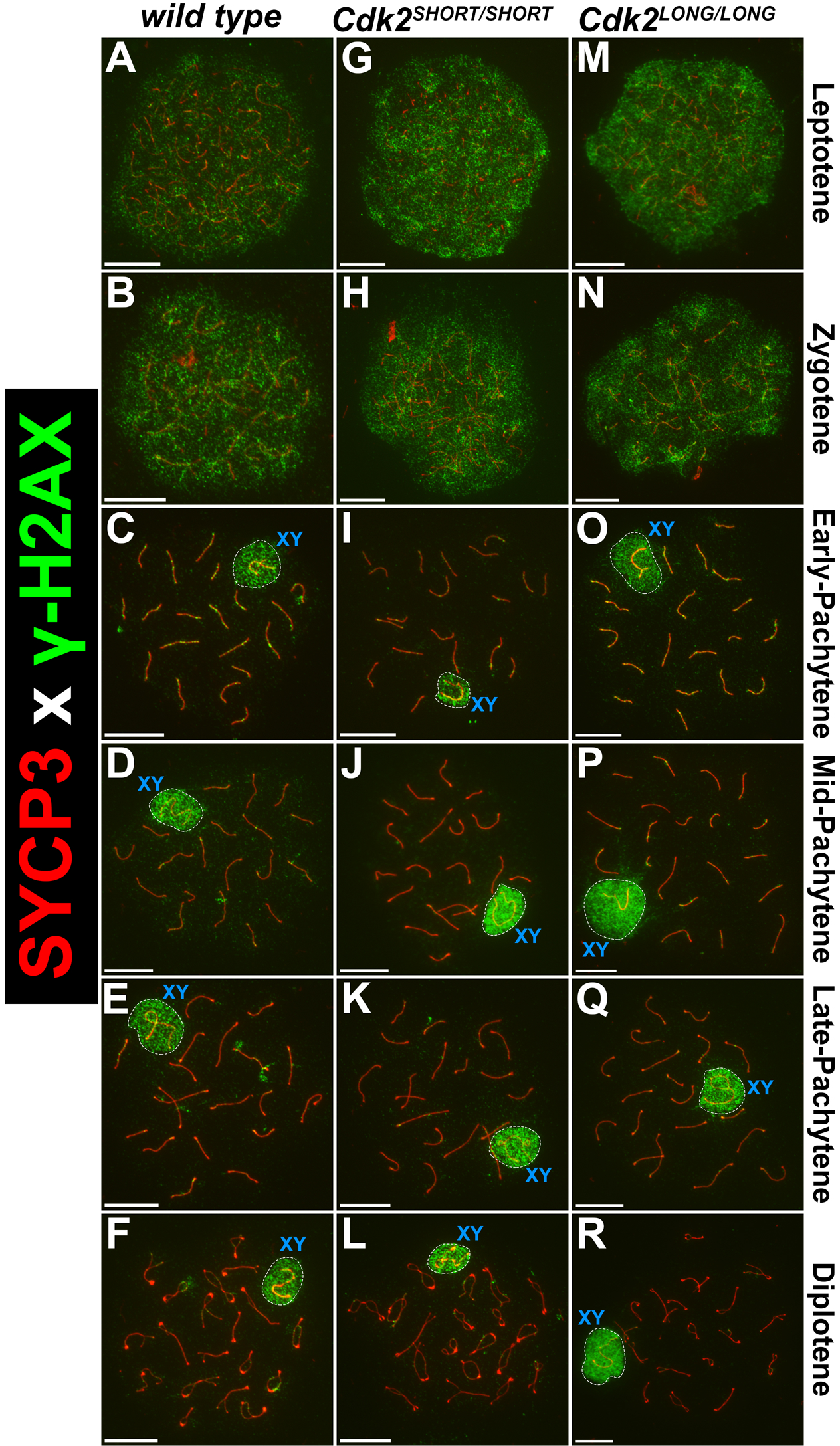
Normal repair of DNA intermediates. Chromosome spreads were prepared from adult testis and stained with antibodies against γH2AX (green, DNA damage) and SYCP3 (red), respectively. Various stages of meiotic prophase I are depicted for wild type (A-F), Cdk2^short/short^ (G-L; CDK2S), and Cdk2^long/long^ (M-R; CDK2L). DNA intermediates are repaired at the correct time with a cloud-like staining in leptotene (A, G, M) and zygotene (B, H, N). Thereafter γH2Ax staining is mostly found along the axis (C, I, O) and retreats over time starting at mid-pachytene (D, J, P, E, K, Q, F, L, R). The sex chromosome are labeled by a blue XY. Scale bars in all panels are 5µm. At least 3 biological replicates were analyzed for each condition.

### Both CDK2L and CDK2S can associate with SpeedyA at the nuclear envelope to maintain normal homolog pairing

The multifunctional nature of CDK2 during meiotic prophase likely arises from the association of CDK2 with a variety of different partner proteins including cyclins, cyclin-like/atypical cyclin proteins [33] and non-cyclins. To date, the contribution of these CDK2-binding partners towards the various actions of CDK2 during meiotic prophase is poorly understood (see review [34]). Currently, one of the most well-characterized meiosis-specific CDK2 partners is the non-cyclin protein, Speedy/Ringo A [35–37]. Like CDK2, Speedy A is required for the pairing of homologous chromosomes and its deletion results in meiotic arrest essentially indistinguishable from that seen upon *Cdk2* deletion [38, 39]. Speedy A has been proposed to promote the loading of CDK2 onto telomeres [39] similarly the deletion of CDK2 leads to a loss of Speedy A binding at telomeres [26]. An added layer of complexity is the observation that the E-type cyclins, especially cyclin E2 are required to maintain the integrity of telomeres during meiotic prophase I [40, 41]. Deletion of Cyclin E2 causes a loss of telomere integrity which also results in pairing defects. This is worsened by the concurrent loss of Cyclin E1 indicating that both E-type cyclins, in addition to Speedy A are required for normal homolog pairing during meiotic prophase. Interestingly, the additional 48 amino acids insertion seen in the CDK2L protein is positioned almost directly opposite the cyclin-binding face of CDK2. This location is distant from the cyclin interface, the ATP-binding, and CKShs1-binding sites [13]. One suggested possibility is that this sequence might influence the interaction between CDK2 and potential binding partners.

CDK2L is known to be expressed preferentially in meiotic cells of both male and female, specifically during the timing of meiotic recombination between homologous chromosomes [39, 42]. The relative expression CDK2S and CDK2L in testis also changes during development. As the levels of CDK2 rise around P14 when the first meiotic cells are observed, the levels of CDK2S fall [22]. Other studies have noted that CDK2L might have preferential binding to Speedy A although this has not been extensively tested. Interestingly in meiotic tissues, the expression pattern of Speedy A mirrors that of CDK2 [22]. Similarly to CDK2L, during meiosis Speedy A can be detected specifically at telomeres and along the sex body [22, 26, 38] but not at late recombination nodules. At the telomeres, SpeedyA has been implicated in recruiting telomere-bound CDK2 to the nuclear envelope to stabilize telomere-nuclear envelope tethering [22].

Base on the previous knowledge, we stained chromosome spreads with antibodies against SpeedyA and pan-CDK2 to detect all isoforms of CDK2. In wild type mice, we found that SpeedyA localizes exclusively to telomeres, whereas CDK2 bind to both telomeres and interstitial sites as was expected (Figure 8; I-IV). At the telomeres, SpeedyA and CDK2 co-localized. In the absence of CDK2L (CDK2^SHORT/SHORT^), SpeedyA binds to telomeres like in the wild type and co-localized with CDK2S at the telomeres but not at the interstitial sites (Figure 8; V-VIII). In the absence of CDK2S in the CDK2^LONG/LONG^ mice, SpeedyA still localized to the telomeres, now overlapping with CDK2L (Figure 8; IX-XII). Therefore, in all three genotypes SpeedyA localization was identical at the telomeres and was thus able to interact with both isoforms of CDK2. Importantly though, the binding partner of CDK2 at the interstitial sites is still unknown.

**Figure 8:**
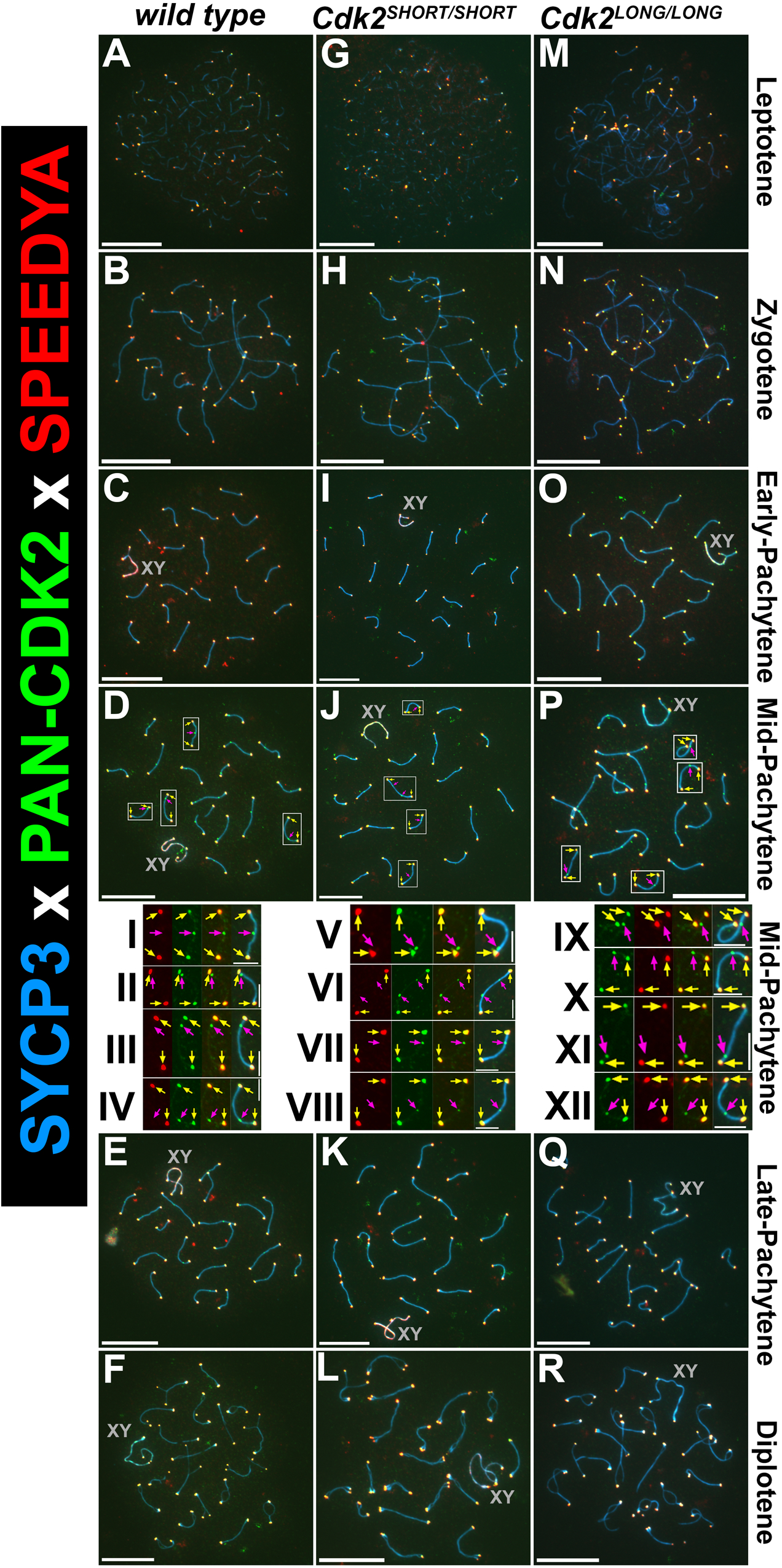
SpeedyA can interact with both CDK2S and CDK2L. Chromosome spreads from adult testis and stained with antibodies against SpeedyA (red), pan-CDK2 (green), and SYCP3 (blue), respectively. Various stages of meiotic prophase I are depicted for wild type (A-F), Cdk2^short/short^ (G-L; CDK2S), and Cdk2^long/long^ (M-R; CDK2L). SpeedyA localizes to the telomeres independent on CDK2 (both CDK2S and CDK2L), CDK2S, or CDK2L. The sex chromosome are labeled by a grey XY. Scale bars in all panels are 5µm; in the insets (I-XII) 1.25µm. At least 3 biological replicates were analyzed for each condition.

### Specific localization of CDK2 in CDK2L and CDK2S mice

As previous studies investigating CDK2 localization during meiosis were done using antibodies detecting both isoforms of CDK2 [25, 26, 32], we designed antibodies to specifically target CDK2L in an attempt to determine whether this isoform displayed specific temporal or special expression during meiotic prophase I. In wild type mice, we found that detection of CDK2 with antibodies specifically against CDK2L detected CDK2 at telomeres but not interstitial sites marking locations of meiotic recombination (Figure 9; I-IV; red). In contrast, pan-CDK2 antibodies that detect both the long and short form of CDK2, stained both telomeres and interstitial sites marking locations of meiotic recombination (Figure 9; I-IV; green). This data indicates that CDK2L has a higher affinity for binding at telomeres compared to interstitial sites. As it is not possible to design specific antibodies against CDK2S which do not cross-react with CDK2L, we determined the binding of CDK2 in our CDK2S mouse model. In the absence of CDK2L (CDK2^SHORT/SHORT^), CDK2S is the only isoform of CDK2 expressed and binds both to the telomeres and the interstitial sites (Figure 9; V-VIII; green). This suggests that CDK2S is able to bind to both sites, at least in the absence of CDK2L. Based on this finding, it was important to check where CDK2L is binding in the absence of CDK2S in the CDK2^LONG/LONG^ mice. Interestingly, CDK2L was localized to both telomeres and interstitial sites, albeit with a stronger signal at the telomeres (Figure 9; IX-XII; red). This suggests that CDK2S and CDK2L can compensate for each other, especially in the absence of the other isoform, which is similar as the compensation seen between different CDKs [24, 43–46].

**Figure 9:**
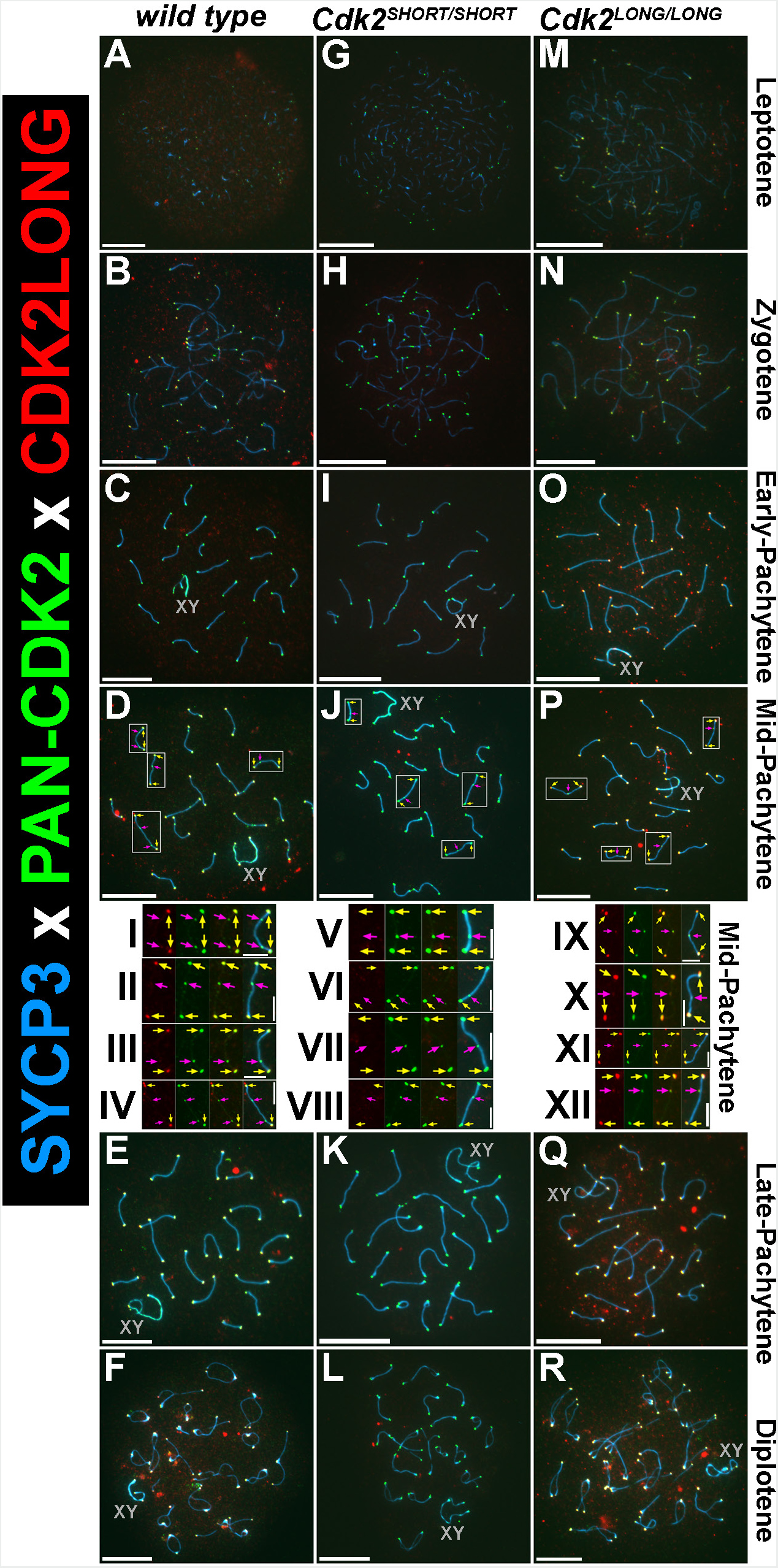
CDK2L can localize to interstitial sites. Chromosome spreads were prepared from adult testis and stained with antibodies against CDK2^LONG^ (red), pan-CDK2 (green; detecting both CDK2S and CDK2L) and SYCP3 (blue), respectively. CDK2S can bind to both telomeres and interstitial sites. The same is true for CDK2L but its affinity to bind to the interstitial sites seems to be lower than CDK2S. The sex chromosome are labeled by a grey XY. Scale bars in all panels are 5µm; in the insets (I-XII) 1.25µm. At least 3 biological replicates were analyzed for each condition.

These results suggest that CDK2L exhibits preferential binding to telomeres but not late recombination nodules during meiotic prophase in wild type mice. Upon sole expression of CDK2S however, the shorter CDK2 isoform can also effectively bind to telomeres, presumably replacing the function of CDK2L. In the converse manner upon the sole expression of CDK2L the longer CDK2 isoform can bind to late recombination nodules in addition to telomeres.

An interesting possibility here is that there exist distinct CDK2-binding proteins at telomeres and late recombination nodules which show preferential binding of CDK2L and CDK2S, respectively. If the preferred CDK2 isoform is absent, however, the remaining pool of CDK2 would be recruited to these sites.

## Discussion

Based on accumulated data from more than 20 years, we set out to investigate the two isoforms of cyclin-dependent kinase 2 (CDK2), CDK2S and CDK2L. In most studies, CDK2S has been investigated but there are a few studies on CDK2L. In order to achieve this, we generated mouse strains that only expressed one of the isoforms but not the other. Using these mice, we studied fertility, which is the major phenotype in *Cdk2^null^* mice, and also cells. Although there might be biochemical differences in CDK2S and CDK2L in vitro, we found that in vivo, these two isoforms can compensate for each other. This resulted in two mouse stains, CDK2S and CDK2L, that are viable and fully fertile.

One of the more interesting and surprising observations, was that although both mouse strains are viable, CDK2L mice seem to be slightly smaller than WT and CDK2S mice. Originally, we and others found that *Cdk2^null^*mice are consistently 10-15% smaller than WT mice [7, 24] but we could not uncover the reasons behind this phenotype. From our current data, it seems that the body weight phenotype is associated either with the expression of CDK2L or the lack of CDK2S. Since also the testis weight followed the same pattern, i.e. decreased in the CDK2L mice, this confirms the overall body weight. Of course since this is a mild phenotype, determining the molecular reason behind the body size decrease will be challenging.

The majority of the efforts reported in this manuscript is dedicated to spermatogenesis, where CDK2 plays multiple roles [47]. Although it was clear that CDK2S and CDK2L mice were fertile, we investigated several stages of meiosis and all the known functions of CDK2 but ultimately had to conclude that CDK2S and CDK2L can compensate for each other’s functions. One of the potential interesting information was that specific antibodies against CDK2L, stained the telomeres very well, as expected, but the staining of the interstitial sites was less pronounced. This confirmed that CDK2L has a high affinity for the telomeres but there was still CDK2 localized to the interstitial sites in the absence of CDK2S (Figure 9, IX-XII). Instead of an effect of affinity, this data could be a result of CDK2 as being part of a complex at the interstitial sites, is less accessible to the antibodies but this does not seem to be the case for the pan-CDK2 antibodies.

Overall, we have found that in vivo both isoforms can perform all the functions that CDK2 is important for. Unfortunately, our work cannot explain why the splice isoforms of CDK2 are conserved in several mammals but not in humans.

The limitations of our study is that despite investing a lot of efforts in probing the known functions of CDK2, there may have been conditions that would have shown a clear difference between the CDK2S and CDK2L mice. Such conditions could include aging, DNA damage in various context like irradiating the mice, or crossing them to *Cdk4^null^* or *p27^Kip1^*mice as we had done with the *Cdk2^null^* mice [11, 43] to determine the compensation by CDK1 and CDK4.

## Materials and methods

### Ethics statement

Experimental procedures were approved by the Animal Care and Use Committee of Biological Resource Centre at Biopolis, A*STAR, Singapore (protocol #171268).

### Generation of Cdk2^Exon6KO^ and Cdk2^Exon6KI^ mice

Cdk2^Exon6KO^ and Cdk2^Exon6KI^ ^mice^ were generated by Cyagen Biosciences (Guangzhou).

### Generation of Exon6KI mice

The gRNA to mouse Cdk2 gene, the donor vector containing Cdk2 gene exon 6 cassette, and Cas9 mRNA were co-injected into fertilized mouse eggs to generate targeted knockin offspring. F0 founder animals were identified by PCR followed by sequence analysis, which were bred to wildtype mice to test germline transmission and F1 animal generation

### gRNAs used for generation of Exon6KI mice

gRNA1 (matching forward strand of gene): AAATGGTATGGAGGCTTGCCAGG gRNA2 (matching forward strand of gene): CCCATTTCCAGGTGACCCGCAGG gRNA3 (matching forward strand of gene): TCTTTGCTGAAATGGTATGGAGG gRNA4 (matching reverse strand of gene): ATAGGGCCCTGCGGGTCACCTGG

Cdk2 gene exon 6 cassette sequence (homology arms are underlined)

<u>AGTACTACTCCACAGCCGTGGATATCTGGAGCCTGGGCTG</u>CATCTTTGCTGAAA TGCACCTAGTGTGTACCCAGCACCATGCTAAGTGCTGTGGGGAACACAGAAGAAATGGAAGACACAGTCTCTGCCCGCTGTGCTCCTATCTAGAAGTGGCTGCATCA CAAGGAGGGGGGATGACCGCAGTGTCTGCCCCACACCCCGTGACCCGCA<u>GGG</u> <u>CCCTATTCCCTGGAGATTCTGAGATTGACCAACTCTT</u>

### Generation of Exon6KO mice

The gRNA to mouse Cdk2 gene, and Cas9 mRNA were co-injected into fertilized mouse eggs to generate targeted knockout offspring. F0 founder animals were identified by PCR followed by sequence analysis, which were bred to wildtype mice to test germline transmission and F1 animal generation.

### gRNAs used for generation of Exon6KO mice

gRNA1 (matches reverse strand of gene): GTTTCGAATAAGAGGTCTATAGG gRNA2 (matches forward strand of gene): ACCGACCCCATGATAAGCCCTGG

### Histology

For hematoxylin and eosin (H&E) staining, testes and ovaries were isolated at the indicated time points and fixed in modified Davidson’s solution, then transferred into 70% ethanol, embedded in paraffin, and cut into 6 mm sagittal sections with a microtome. To evaluate proliferation in the ovary, histological sections were stained using Ki-67 antibodies (Leica Microsystems, NCL-Ki67p). Images were captured using either an Olympus BX61 or Zeiss Imager Z1 microscope.

### Western blots

The preparations of protein extracts and western blotting procedures were done as described [30].

### Meiotic chromosome spreads from M.musculus testes

This procedure was done as we have described in detail previously [26].

## Acknowledgements

We thank all the past and present members of the Kaldis lab for their support and input.

PK is supported by the Swedish Research Council (2021-01331), the Swedish Cancer Society (Cancerfonden; 21-1566Pj), the Crafoord Foundation (Ref. No. 20220628), the Faculty of Medicine, Lund University, the Swedish Foundation for Strategic Research Dnr IRC15-0067, and Swedish Research Council, Strategic Research Area EXODIAB, Dnr 2009-1039. KL was supported by a grant from the Shenzhen Science and Technology Program (KQTD20190929172749226) China. The funders had no role in study design, data collection and interpretation, or the decision to submit this manuscript for publication.

## Author Contributions

Conceptualization: Nathan Palmer, Philipp Kaldis

Data curation: Nathan Palmer, S. Zakiah A. Talib, Christine M.F. Goh, Jin Rong Ow, Tommaso Tabaglio

Formal analysis: Nathan Palmer

Funding acquisition: Kui Liu, Philipp Kaldis

Investigation: Nathan Palmer, S. Zakiah A. Talib, Christine M.F. Goh, Jin Rong Ow, Tommaso Tabaglio

Methodology: Nathan Palmer, S. Zakiah A. Talib, Christine M.F. Goh, Jin Rong Ow, Tommaso Tabaglio

Project administration: Philipp Kaldis Resources: Kui Liu, Philipp Kaldis

Supervision: Ernesto Guccione, Philipp Kaldis

Validation: Nathan Palmer, S. Zakiah A. Talib, Christine M.F. Goh, Jin Rong Ow, Tommaso Tabaglio

Writing – original draft: Nathan Palmer, Philipp Kaldis

Writing – review & editing: Nathan Palmer, Ernesto Guccione, Philipp Kaldis (with the input of all authors)

